# Very long chain fatty acids drive 1-deoxy-Sphingolipid toxicity

**DOI:** 10.1101/2025.05.13.653734

**Authors:** Adam Majcher, Gergely Karsai, Elkhan Yusifov, Martina Schaettin, Ermanno Malagola, Peter Horvath, Jinmei Li, Maftuna Shamshiddinova, Gai Zhibo, Raghvendra Dubey, Tim Peterson, Sofía Rodriguez-Gallardo, Kuniyoshi Shimizu, Takeshi Harayama, Thorsten Hornemann

## Abstract

1-deoxy-sphingolipids (1-deoxySLs) are atypical sphingolipids synthesized by the serine palmitoyltransferase (SPT) when L-alanine is used instead of its canonical substrate L-serine. Increased 1-deoxySLs are associated with sensory neuropathies such as Hereditary Sensory and Autonomic Neuropathy type 1 (HSAN1) and diabetic polyneuropathy (DPN). Despite their known cellular, mitochondrial, and neurotoxic effects, the mechanisms underlying their toxicity remain poorly understood.

Using a CRISPR interference (CRISPRi) screening approach, we identified CERS2, ELOVL1, ACACA, HSD17B12, and PTPLB as key mediators of 1-deoxySL-induced toxicity. All genes are integral to the biosynthesis of very long-chain (VLC) fatty acids and VLC-ceramides. We validated these findings through genetic knockdown experiments, cytotoxicity assays, and stable isotope-resolved lipidomics via LC-MS/MS. Pharmacological inhibition of ELOVL1 using a preclinical tested compound alleviated the cellular, mitochondrial, and neuronal toxicity induced by 1-deoxySLs.

Supplementation experiments combining 1-deoxySLs with various VLC fatty acids revealed that 1-deoxyDHceramide conjugated to nervonic acid (m18:0/24:1) is the principal toxic specie. Further mechanistic studies showed that m18:0/24:1 induces apoptosis through the mitochondrial permeability transition pore (mPTP) formation. Inhibition of BAX or blocking mPTP formation with cyclosporin A effectively prevented toxicity.

In conclusion, our findings demonstrate that 1-deoxyDHCeramides conjugated to nervonic acid are the primary mediators of 1-deoxySL toxicity, acting through mitochondrial dysfunction and BAX-dependent apoptosis.

## Introduction

Sphingolipids (SL) are a structurally diverse group of bioactive lipids that share a a long-chain base (LCB) as common core structure. Forming the LCB is the rate limiting reaction in the SL *de novo* synthesis and initiates with the conjugation of L-Serine and Palmitoyl-CoA, catalyzed by the Serine-Palmitoyltransferase (SPT). The resulting product 3-ketosphinganine is rapidly converted by ketodihydrosphingosine reductase (KDSR) to Sphinganine (Sa, d18:0), which serves as a substrate for a group of Ceramide synthases (CerS) that N-acylate the LCB with a long chain (LC) - or very long chain (VLC) fatty acid. Mammalians express six CERS isoforms (CERS1-6) with different specificities towards the length and saturation state of their fatty acid substrates. The product, dihydroCeramide (DHCer) is finally converted to Ceramides (Cer) by the Dihydroceramide desaturase 1 (DEGS1), by introducing a Δ4E double bond into the LCB backbone. Ceramides are the central molecules of SL metabolism and serve as building blocks for complex SL, such as Sphingomyelins and Glycosphingolipids ^1^.

In addition to L-Serine, SPT can also utilize L-Alanine forming the atypical LCB 1-deoxysphinganine (1-deoxySa, m18:0) which in comparison to SA lacks the essential hydroxyl group at C1 position ^2^. Like SA also 1-deoxySa is metabolized by CERS forming 1-deoxy-dihydroCeramides (1-deoxyDHCer, m18:0/xx:y) ^3^, but in contrast to canonical SL, 1-deoxyDHCer are not further metabolized by DEGS1 but instead by fatty acyl desaturase 3 (FADS3) which introduces a Δ14Z, instead of a Δ4E double bond ^4^.

As 1-deoxySL lack the C1-hydroxylic group of canonical LCBs, they cannot get converted into complex SL, but also not degraded via the canonical SL catabolism, which requires the formation of the catabolic intermediate Sphingosine-1-Phosphate (S1P) ^3^.

1-deoxySL are toxic to neurons and other cells ^4-8^, and have been linked to peripheral sensory neuropathies such as the Hereditary Sensory Neuropathy Type 1 (HSAN1)^9-12^, diabetic neuropathy^13-16^ as well as the rare retinopathy Macular Telangiectasia type 1 (MacTel)^17^. 1-deoxySLs have been shown to trigger various pathological cellular processes, such as altered mitochondrial fission, increased radical oxygen species (ROS), autophagy, endoplasmic reticulum (ER) stress, unfolded protein response, cytoskeletal defects, and calcium handling abnormalities ^5,6,8,18-21^.

Despite these findings, the toxicity of 1-deoxySL and the underlying molecular mechanisms are not yet understood. In this study, we combine a CRISPRi-based functional genomics screen, with functional lipidomics, targeted genetic interference and pharmacological studies, to understand the metabolic and structural determinants of 1-deoxySL toxicity.

## Results

### 1-deoxyDHCeramides are the primary mediators of 1-deoxySphingolipids induced toxicity

To investigate the molecular mediators of 1-deoxysphingolipid (1-deoxySL) toxicity, we employed the ceramide synthase (CERS) inhibitor Fumonisin B1 (FB1), a mycotoxin previously reported to mitigate 1-deoxySa-induced cytotoxicity (Figure 1 a) ^20,22^. Co-treatment of HeLa cells with 1-deoxySa and FB1 led to a significant reduction in toxicity, confirming that CERS inhibition has a protective effect (Figure 1b). To investigate the underlying metabolic changes, we conducted a stable isotope-labeled flux assay by supplementing _d3_-1-deoxySa for 24 hours, followed by high-resolution LC-MS/MS-based lipidomics to trace the incorporation of the labeled LCB into downstream sphingolipid species. FB1 treatment led to a marked accumulation of free long-chain bases (LCBs), including both d_3_-1-deoxySa and d_3_-1-deoxySo, while simultaneously suppressing the formation of N-acylated products, including d_3_-1-deoxyDHCer and d_3_-1-deoxyCer (Figure 1c,d). These findings suggest that toxicity is not mediated by the LCBs themselves, but rather by their N-acylated derivatives.

**Figure 1.**
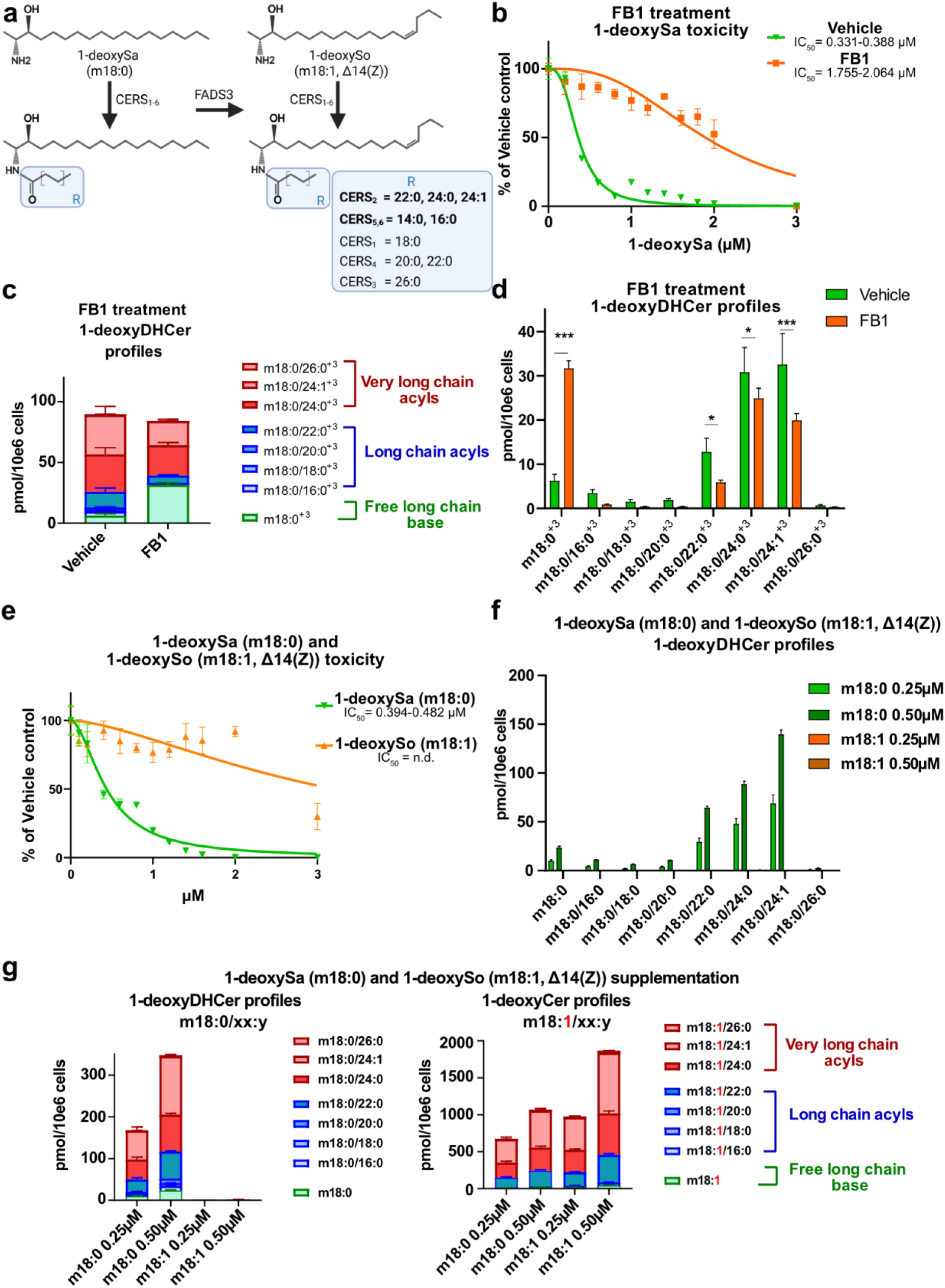
1-deoxyDHCeramides are the primary mediators of 1-deoxySphingolipids induced toxicity. **a** Schematic representation of 1-deoxySL metabolism. The horizontal arrow indicates desaturation of 1-deoxysphinganine (1-deoxySa, m18:0) at the Δ14(Z) position catalyzed by fatty acid desaturase 3 (FADS3). Vertical arrows represent N-acylation of 1-deoxySa by ceramide synthase isoforms (CERS1–6). The blue box highlights the acyl-CoA substrate specificities of individual CERS isoforms. **b** 1-deoxySa (m18:0) toxicity in HeLa cells in the presence or absence of the pan-CERS inhibitor fumonisin B1 (FB1, 35 µM). Toxicity was determined by CellTiter-GLO assay as described in the methods section. **c, d** Formation of d3-1-deoxyDHCer (m18:0/xx:y^+3^) species in HeLa cells supplemented with d3-1-deoxySa and treated with pan-Ceramide synthase inhibitor Fuminosin B1 (FB1, 35µM) or Vehicle. **e** Comparison of cytotoxicity of 1-deoxySa or (m18:0) 1-deoxysphingosine (1-deoxySo, m18:1, Δ14(Z)) on HeLa cells. **f, g** Intracellular accumulation of 1-deoxyDHCer and 1-deoxyCer species formed in HeLa cells supplemented with 1-deoxySa (m18:0) or 1-deoxySo (m18:1, Δ14(Z)). All lipid data were measured by an untargeted high resolution LC-MS/MS lipidomics workflow and normalized to an internal standard and cell count. Data are shown as mean ± SD. Toxicity data were normalized to vehicle treated cells and represented as mean ± SD (N=4). Non-linear regression analysis was used to calculate toxicity curves and IC50 values (95% Confidence Interval).

Next, we investigated whether the presence of the Δ14Z double bond has an influence on toxicity. To assess the role of the Δ14(Z) double bond introduced by fatty acid desaturase 3 (FADS3, Figure 1a), we compared the cytotoxicity of 1-deoxySa (saturated, m18:0) and its unsaturated analog 1-deoxySo (m18:1, Δ14(Z)). Consistent with previous reports implicating FADS3 in detoxification^4^, the addition of 1-deoxySa exhibited significantly higher cytotoxicity compared to 1-deoxySo (Figure 1 e). To determine the metabolic fate of these LCBs, we performed an untargeted lipidomics of HeLa cells supplemented with either 1-deoxySa (m18:0) or unsaturated 1-deoxySo (m18:1, Δ14(Z)). Supplementation with 1-deoxySa resulted in the accumulation of both 1-deoxyDHCer and 1-deoxyCer species, whereas 1-deoxySo was metabolized exclusively to 1-deoxyCer (Figure 1f,g). Together, these data indicate that saturated N-acylated species, particularly 1-deoxyDHCer, are the principal mediators of 1-deoxySL-induced cytotoxicity.

### CRISPRi toxicity screen identifies gene candidates responsible for 1-deoxySphingolipids toxicity

To identify genes involved in in 1-deoxySL toxicity, we performed a systematic CRISPRi toxicity screen. Haploid K562 cells expressing CRISPRi-dCas9-KRAB were transfected with a CRISPRi v2 sgRNA library and cultured for five passages in presence of 1-deoxySa (1.5µM) which corresponds to the experimentally determined IC_50_ under the screening conditions (Figure 2 a). Comparative analysis of sgRNA abundance revealed 5 genes whose knockdown conferred a significant resistance to 1-deoxySa induced toxicity (Figure 2b). All identified genes are functionally linked to the elongation of fatty acids (ACACA, HSD17B12, PTPLB), the synthesis of VLC-FAs (ELOVL1) or the incorporation of VLC-FAs into the LCB backbones (CERS2) (Figure 2c). Notably, ELOVL1 catalyzes the elongation of saturated and monounsaturated acyl-CoAs to VLC-FAs such as 22:0, 24:0, and 24:1, which are the substrates for CERS2, forming VLC-1-deoxyDHCer species^23^. These findings position ELOVL1 directly upstream of CERS2 in the biosynthetic VLC-FA pathway. Conversely, FADS3 was identified as a sensitizing gene, consistent with the protective effect of the Δ14(Z) double bond observed in earlier experiments (Figure 1e). To further assess the functional context of these hits, we performed gene set enrichment analysis (GSEA) using logFC values derived from the CRISPRi screen. This revealed significant enrichment of genes involved in the biosynthesis of unsaturated fatty acids and elongation among those promoting toxicity (Figure 2d). In contrast, protective gene signatures were more broadly distributed across pathways related to ribosomal function, oxidative phosphorylation, DNA replication, RNA transport, and peroxisomal activity (Figure 2d). This highlights the central role of the lipid metabolic networks, particularly VLC-FA biosynthesis, in mediating 1-deoxySL-induced cytotoxicity.

**Figure 2.**
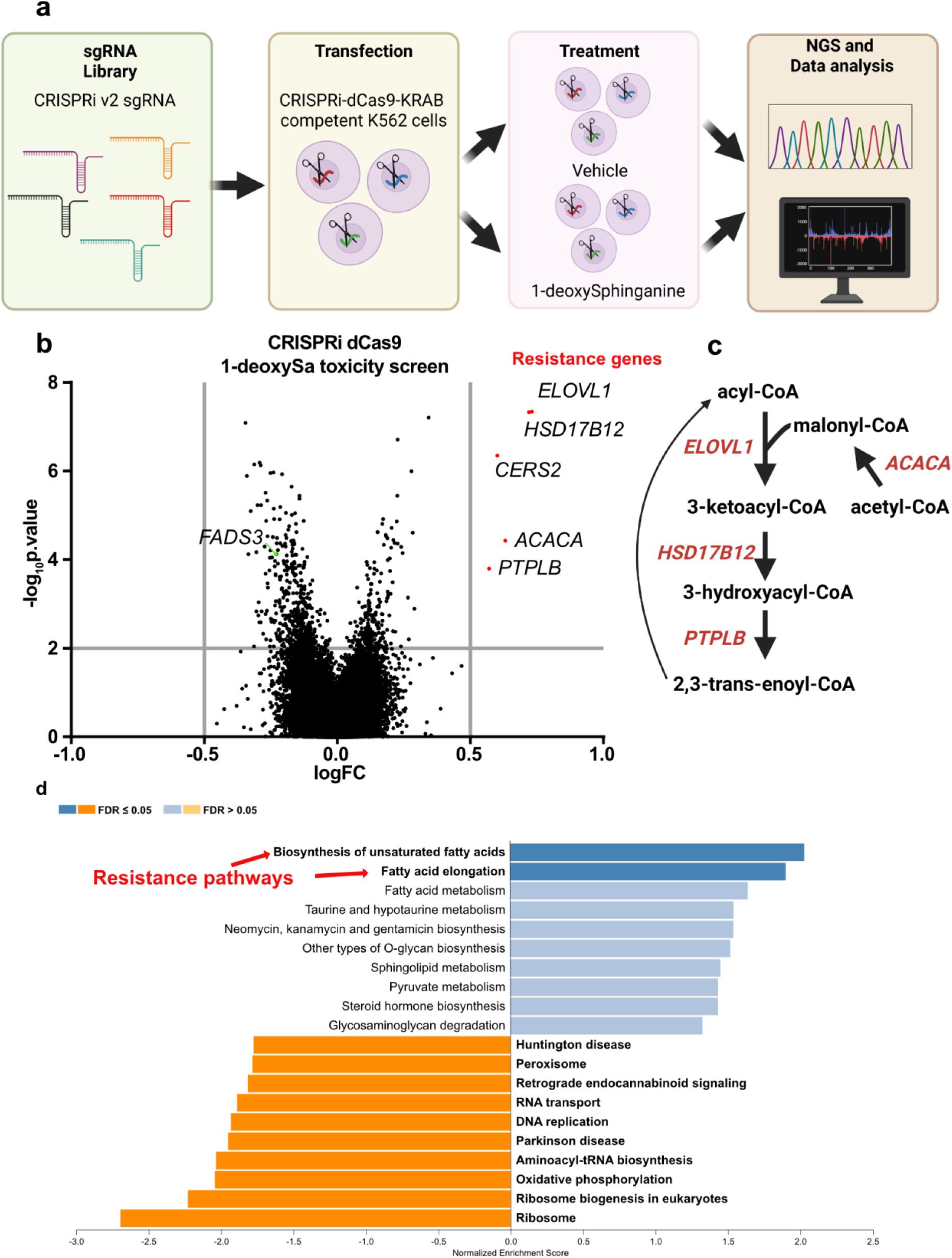
CRISPRi toxicity screen identifies gene candidates responsible for 1-deoxySphingolipids toxicity. **a** Schematic representation of the CRISPRi-based toxicity screen. Haploid K562 CRISPRi-dCas9-KRAB cells were transduced with the CRISPRi v2 sgRNA library and cultured for five passages in the presence of 1-deoxySa (1.5µM) or vehicle control. **b** Volcano plot displaying differentially enriched genes in 1-deoxySa-treated cells. The log fold change (logFC) indicates the difference in sgRNA as quantified by RNA sequencing. Genes conferring resistance to 1-deoxySa toxicity (logFC ≥ 0.5, q-value ≤ 0.01) are shown in red. **c** Schematic overview of the fatty acid elongation pathway showing the position of key genes identified in the screen. ELOVL1, HSD17B12, ACACA, and PTPLB are involved in very-long-chain fatty acid (VLC-FA) biosynthesis, while CERS2 incorporates VLC-FAs into ceramide and 1-deoxyDHCer species. Gene set enrichment analysis (GSEA) of logFC values derived from the CRISPRi screen. Pathways were assigned using the KEGG database. Genes sensitizing cells to 1-deoxySa clustered within unsaturated fatty acid biosynthesis and elongation pathways, whereas protective genes were associated with diverse pathways including ribosome biogenesis, oxidative phosphorylation, DNA replication, RNA transport, and peroxisomal function.

### Silencing of ELOVL1 and CERS2 expression confirms their role in 1-deoxysphingolipid mediated toxicity

Based on the CRISPRi screen, we focused on the functional validation of ELOVL1 and CERS2 as key mediators of 1-deoxySL toxicity (Figure 3a).

**Figure 3.**
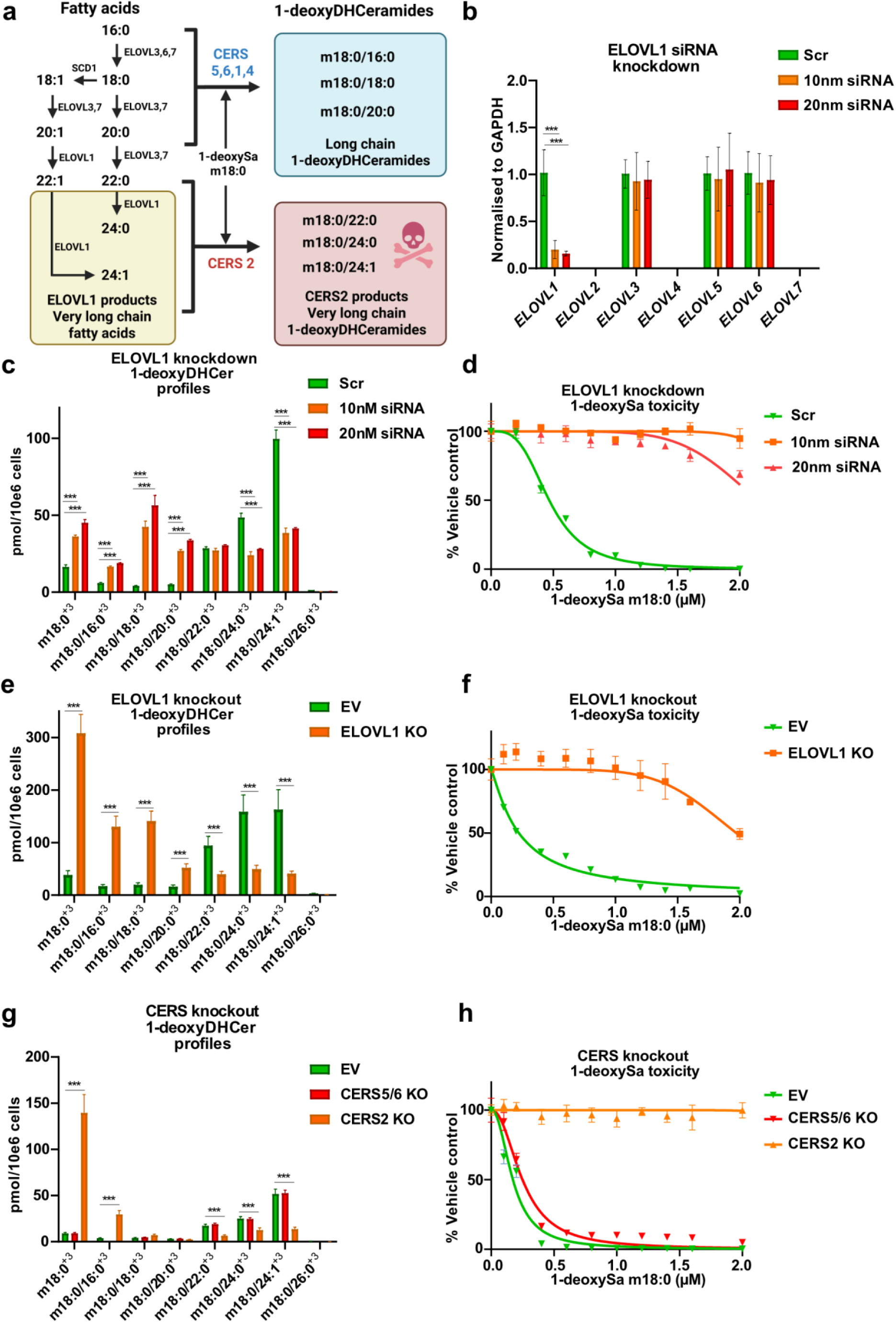
Genetic targeting of ELOVL1 and CERS2 confirms their role in 1-deoxysphingolipids mediated toxicity. **a** Schematic overview of the metabolic pathway linking fatty acid elongation by ELOVL1-7 to 1-deoxyDHCeramide synthesis by ceramide synthases (CERS1-6). **b** mRNA expression levels of ELOVL1–7 in HeLa cells following siRNA-mediated knockdown of ELOVL1. Silencing ELOV1 did not affect expression of other ELOVL isoforms. Expression levels were determined using RT-qPCR and normalized to GAPDH. **c** Quantification of isotopically labeled 1**-**deoxyDHCer (m18:0/xx:y)^+3^ species in HeLa cells after siRNA ELOVL1 knockdown and subsequent d_3_-1-deoxySa supplementation. **d** Toxicity of 1-deoxySa in HeLa cells transfected with scrambled or ELOVL1-targeting siRNAs (10 or 20 nM), 72 h prior to treatment (n = 3). **e** Quantification of isotopically labeled 1-deoxyDHCer species (m18:0/xx:y^+3^) in CRISPR Cas9 generated ELOVL1 KO and control HeLa cells (EV, Empty vector treated) after d_3_-1-deoxySa supplementation **f** 1-deoxySa-induced toxicity in ELOVL1 KO and control HeLa cells (EV). **g** Abundance of d₃-1-deoxyDHCer species (m18:0/xx:y^+3^) in CRISPR Cas9 CERS2 KO, CERS5/6 KO, and EV control HeLa cells following d₃-1-deoxySa supplementation. **h** Toxicity of 1-deoxySa in CERS2KO, CERS5/6KO and Control (EV) cells. Lipid species were quantified by untargeted high-resolution LC-MS/MS and normalized to internal standards and cell counts. Bar graphs are presented as mean ± SD (n = 3). Toxicity data were measured by the CellTiter-GLO assay and normalized to vehicle-treated controls and shown as mean ± SD (n = 4). Significance was determined using two-tailed unpaired Student’s t-test with multiple testing correction (two-stage step-up method of Benjamini, Krieger, and Yekutieli). *p < 0.05, **p < 0.01, ***p < 0.001. IC_50_ values were calculated by nonlinear regression with 95% confidence intervals.

To assess the role of ELOVL1, we performed siRNA-mediated knockdown in HeLa cells. RT-qPCR confirmed selective silencing of ELOVL1, with no off-target effects on other ELOVL isoforms (Figure 3b). To evaluate the metabolic consequences, we conducted a d_3_-1-deoxySa isotopic flux assay coupled to high-resolution LC-MS/MS lipidomics. ELOVL1 knockdown led to a significant reduction in very-long-chain (VLC) 1-deoxyDHCer species, specifically in m18:0/24:0 and m18:0/24:1, and a compensatory increase in long-chain (LC) 1-deoxyDHCer species (m18:0/16:0, m18:0/18:0, m18:0/20:0) (Figure 3c). This shift in sphingolipid composition was accompanied by a significant reduction in 1-deoxySa induced cytotoxicity (Figure 3d).

To further confirm these findings, we used CRISPR-Cas9-generated ELOVL1 knockout (KO) HeLa cells. As expected, ELOVL1 KO resulted in decreased flux of d_3_-1-deoxySa into VLC 1-deoxyDHCer species, with a corresponding increase in LC 1-deoxyDHCer species, recapitulating the knockdown phenotype (Figure 3e). These data establish ELOVL1 as a critical upstream enzyme modulating the accumulation of cytotoxic VLC 1-deoxyDHCer species.

We next examined the roles of the CERS2 and CERS5/6 isoforms. While CERS2 primarily utilizes VLC acyl-CoAs (22:0, 24:0, 24:1), CERS5 and CERS6 preferentially process LC fatty acyl-CoAs (16:0, 18:0). To investigate their functions, we used CRISPR-Cas9 to generate CERS2 knockout (KO) and CERS5/6 double KO HeLa cell lines, which were compared to empty vector (EV) controls.

CERS2 KO cells showed a substantial reduction in d_3_-1-deoxySa-derived VLC 1-deoxyDHCer species (m18:0/22:0, m18:0/24:0, m18:0/24:1), along with elevated levels of LC 1-deoxyDHCer and free LCB 1-deoxySa (Figure 3g). In contrast, CERS5/6 KO cells had a reduced formation of LC 1-deoxyDHCer species, with minimal effects on VLC species. Functional assays confirmed that absence of CERS2 conferred significant protection against 1-deoxySa toxicity, whereas CERS5/6 KO cells had only a marginal influence on the 1-deoxySL induced toxicity (Figure 3h).

Together, these results validate ELOVL1 and CERS2 as key enzymes driving the synthesis of cytotoxic VLC 1-deoxyDHCer species and mediating 1-deoxySL-induced cell death.

### Pharmacological inhibition of ELOVL1 mitigates 1-deoxysphingolipids induced mitochondrial and neuronal toxicity

To evaluate whether a pharmacological inhibition of ELOVL1 could serve as potential therapeutic strategy to reduce 1-deoxySL toxicity, we employed the previously reported small-molecule inhibitor Compound 22 (22) (CAS 2761063-99-2). The pyrimidine-ether compound has been shown to be biologically active in mouse and rat models, with established repeat-dose toxicology studies in cynomolgus monkeys ^24,25^. This compound was originally developed as a potent and selective ELOVL1 inhibitor in the context of X-linked adrenoleukodystrophy (X-ALD) with well characterized in vitro and in vivo activities (Figure 4a) ^24,25^.

**Figure 4.**
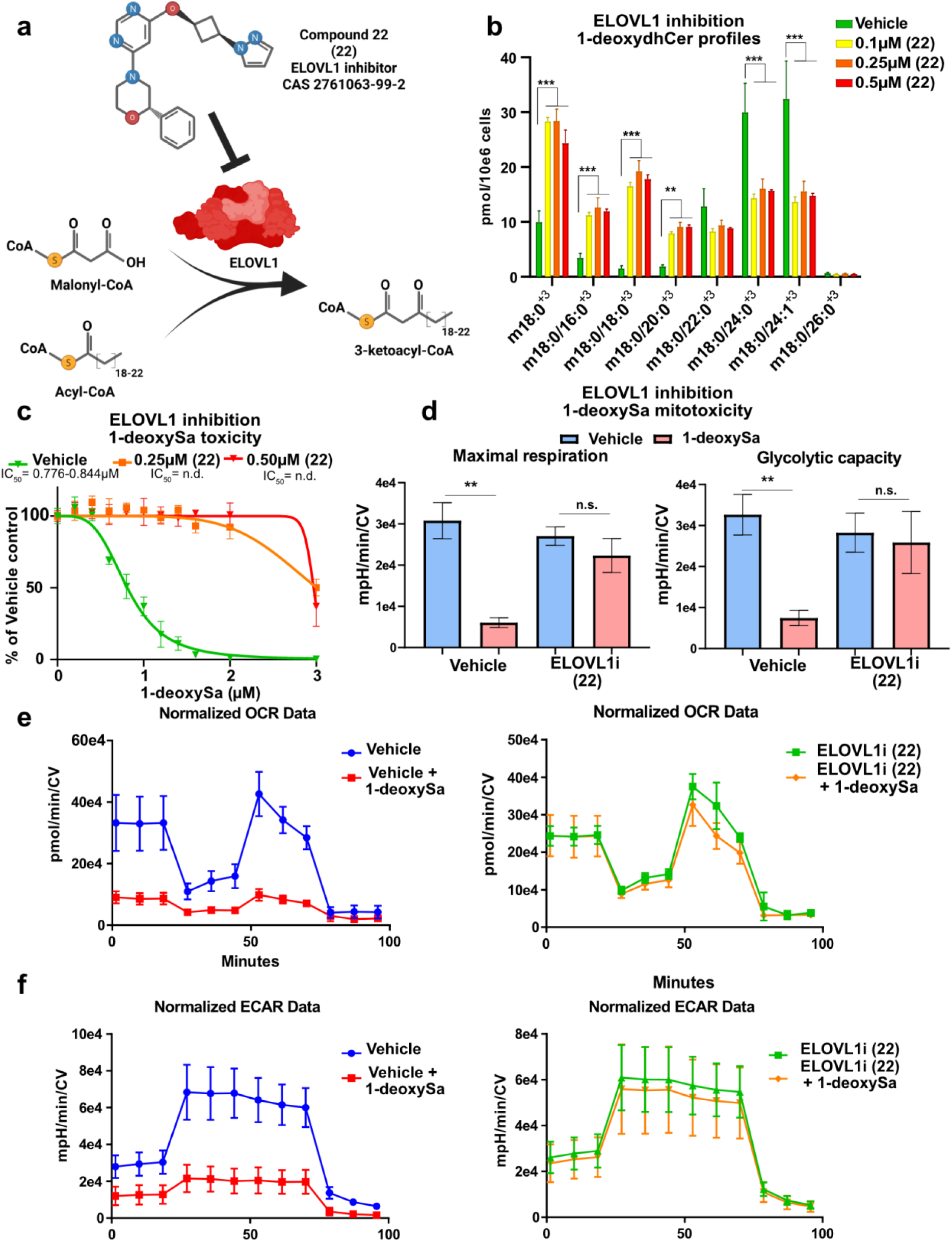
Pharmacological inhibition of ELOVL1 mitigates 1-deoxysphingolipids induced mitochondrial and neuronal toxicity. **a** Schematic enzymatic reaction catalyzed by ELOVL1 and its inhibition by the small-molecule inhibitor Compound 22 (ELOVL1i; (22); CAS 2761063-99-2).**b** Quantification of 1-deoxyDHCer (m18:0/xx:y^+3^) species in HeLa cells supplemented with d_3_-1-deoxySa and ELOVL1 inhibitor (22) or vehicle control. **c** Cytotoxicity of 1-deoxySa in HeLa cells treated with ELOVL1i (0.25 µM) or vehicle. Data were normalized to vehicle-treated controls and shown as mean ± SD (n = 4). IC_50_ values were derived from nonlinear regression analysis with 95% confidence intervals.**d** Maximal respiration and glycolytic capacity of HeLa cells treated with vehicle, 1-deoxySa (0.5 µM), or 1-deoxySa + ELOVL1i (0.25 µM), measured using a Seahorse XF analyzer. **e** Oxygen consumption rate (OCR) and **f** Extracellular acidification rate (ECAR) over time in vehicle- or ELOVL1i (22)-treated HeLa cells following exposure to 1-deoxySa (0.5 µM) measured using a Seahorse XF analyzer.All Seahorse data were normalized to crystal violet staining (CV) performed after the assay to correct for cell number and viability. This normalization was essential, as 1-deoxySa-induced cytotoxicity significantly affects viable cell count and could confound metabolic measurements. Data are shown as mean ± SD (N=4).

In HeLa cells, treatment with (22) significantly reduced the metabolic flux of d_3_-1-deoxySa into very-long-chain (VLC) 1-deoxyDHCer species (m18:0/24:0 and m18:0/24:1), and again caused a compensatory increase in LC-1-deoxyDHCer species (m18:0/16:0, m18:0/18:0, m18:0/20:0) (Figure 4b), resulting in a significant protection from 1-deoxySa-induced cytotoxicity (Figure 4c).

Previous studies have linked elevated levels of 1-deoxySLs to mitochondrial dysfunction and structural damage, including mitochondrial fragmentation ^26^. We therefore tested whether compound 22 could prevent 1-deoxySL-induced mitochondrial damage. We performed a Seahorse XF Mito Tox assays in HeLa cells, treated with 1-deoxySa (0.5 µM). We observed a marked reduction in mitochondrial respiration (OCR), glycolytic capacity (ECAR), and maximal respiratory function. A co-treatment with (22) fully restored all functional parameters (Figure 4d, e, f). All mitochondrial data were normalized to crystal violet staining to account for cell loss after 1-deoxySa treatment and enable accurate assessment of mitochondrial activity.

To confirm the metabolic and protective effects of (22) in a physiological neuronal context, we established an in vivo flux model based on a live chicken embryo model. Living chicken embryos were injected with d_3_-1-deoxySa ± (22) as described, followed by dissection of dorsal root ganglia (DRG) and high-resolution LC-MS/MS lipidomics profiling (Extended Figure 4a). Treatment with (22) significantly reduced the accumulation of VLC 1-deoxyDHCer species (m18:0/22:0, m18:0/23:0, m18:0/24:0, m18:0/24:1) and induced a compensatory increase in long-chain (LC) species (m18:0/16:0, m18:0/18:0) (Extended Figure 4b). These results demonstrate that (22) is metabolically active in the nervous system, particularly in sensory neurons, which are the primary targets of 1-deoxySL toxicity. However, due to technical limitations, the effects on the developing peripheral nervous system of the embryo have not yet been evaluated. Instead, we assessed the neuroprotective potential of (22), in dissociated chicken DRG neurons obtained from day-9 chicken embryos by supplementing 1-deoxySa (0.05 or 0.1 µM) in presence or absence of (22). Neurons were stained for neurofilament-M and neurite area was quantified by fluorescence microscopy and image analysis (Extended Figure 4c). 1-deoxySa treatment caused a significant loss of neurites which was effectively rescued by (22) (Extended Figure 4d, e). Notably, (22) alone had no adverse effect on neurite morphology, indicating that ELOVL1 inhibition itself is not intrinsically neurotoxic. Comparable neuroprotective effects were observed in differentiated SH-SY5Y neuroblastoma cells, where (22) prevented neurite collapse following 1-deoxySa exposure (data not shown).

**Extended Figure 4.**
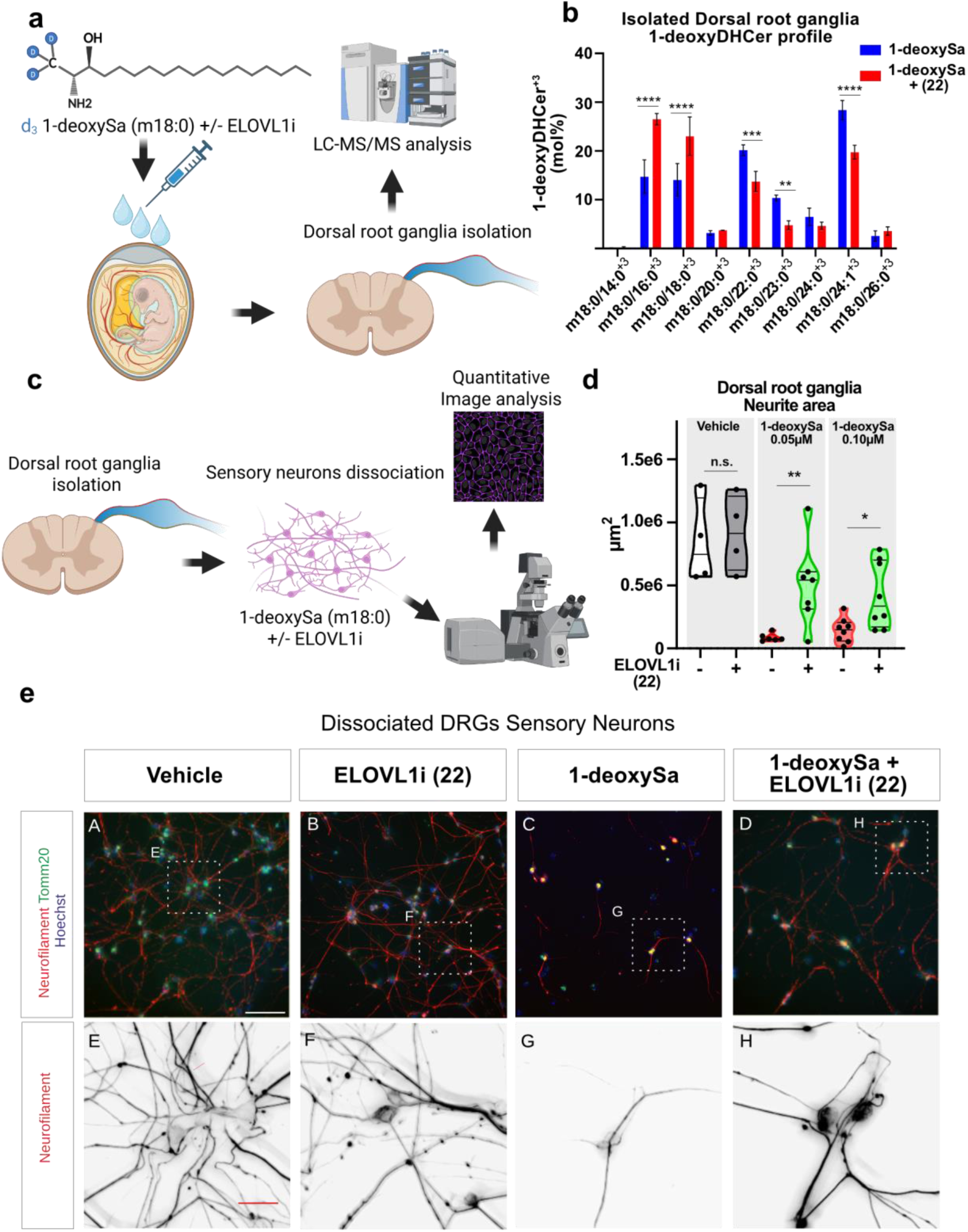
Pharmacological inhibition of ELOVL1 mitigates 1-deoxysphingolipids induced mitochondrial and neuronal toxicity. **a** Schematic overview of the in vivo d_3_-1-deoxySa (m18:0) flux assay using whole chicken embryos. Embryos were supplemented with d_3_-1-deoxySa (m18:0), followed by isolation of dorsal root ganglia (DRG) and high-resolution LC-MS/MS lipidomics analysis. **b** Quantification of 1-deoxyDHCer (m18:0/xx:y^+3^) species in DRG after the whole chicken embryo supplementation with d_3_-1-deoxySa in presence or absence of the ELOVLi (22). Data were normalized to internal standards and sum of all labeled 1-deoxyDHCer. **c** Schematic of the DRG neurotoxicity assay. DRGs were isolated from 9-day-old chicken embryos, dissociated, and cultured with 1-deoxySa in the presence or absence of the ELOVL1 inhibitor (22). Neuronal toxicity was assessed via fluorescence microscopy and quantitative image analysis. **d** Quantification of neurite area in dissociated DRG neurons treated with 1-deoxySa (0.05 µM or 0.1 µM) or vehicle (DMSO), in the presence or absence of ELOVL1i (22). Neurites were labeled with anti-Neurofilament-M, mitochondria with anti-TOMM20, and nuclei with DAPI. Imaging was performed at 20× magnification, and total neurite area was quantified using ImageJ.**e** Representative fluorescence images of DRG neurons treated with vehicle or 1-deoxySa (0.05 µM) ± ELOVL1i (22). Panels **A–D** show axonal networks at 20× magnification; panels **E–H** show close up images along axons. Scale bars: 100 µm (**A–D**); 25 µm (**E–H**).

In summary, these findings demonstrate that compound (22), a preclinical validated pharmacological ELOVL1 inhibitor, protects against 1-deoxySL-induced mitochondrial dysfunction and neurotoxicity, supporting its potential as a therapeutic strategy for disorders involving pathological 1-deoxySL accumulation.

### Nervonyl-1-deoxyDHCeramide (m18:0/24:1) induces mitochondrial damage via mPTP and BAX, revealing a novel mechanism of 1-deoxysphingolipid cytotoxicity

Finally, we aimed to see whether toxicity is different for the individual LC and VLC 1-deoxyDHCer species (Figure 5 a). ELOVL1 inhibition prevents the formation of VLC 1-deoxyDHCer synthesis but does not directly inhibit CERS2 activity. Therefore we aimed to resupplement individual fatty acid substrates (16:0, 24:0 and 24:1) in presence of (22) to address the toxicity of the individual 1-deoxyDHCer species. As VLC-FA’s are typically not absorbed efficiently by the cells, we developed a protocol that allowed an efficient resorption of the supplemented FAs. For that, FAs were supplemented as Na+ salts in complex with bovine serum albumin (BSA) (methods section) and added in presence of (22) to block further elongation.

**Figure 5.**
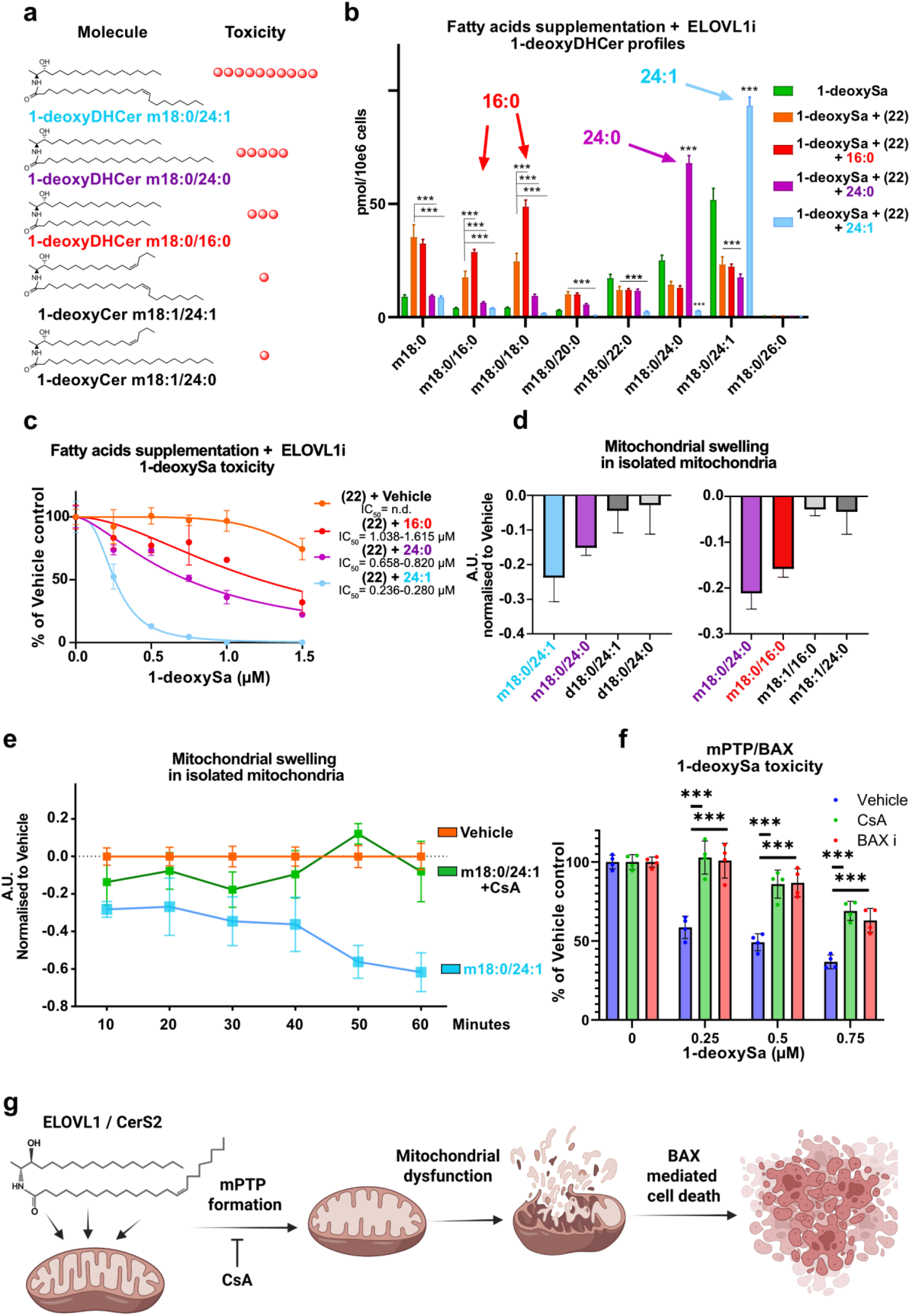
Nervonyl-1-deoxyDHCeramide (m18:0/24:1) induces mitochondrial damage via mPTP and BAX, revealing a novel mechanism of 1-deoxysphingolipid cytotoxicity. **a** Schematic representation of the chemical structures of individual 1-deoxysphingolipid (1-deoxySL) species and their relative cytotoxic potential.**b** Quantification of 1-deoxyDHCer species in HeLa cells supplemented with 1-deoxySa and co-treated with long-chain (LCFA, 25µM) or very-long-chain fatty acids (VLCFA, 25 µM), in presence of ELOVL1 inhibitor Compound (22) (0.25 µM). Fatty acids were applied as sodium salts complexed with 4 mg/mL bovine serum albumin (BSA); BSA alone served as vehicle control. Data are shown as mean ± SD (n = 3).**c** 1-deoxySa cytotoxicity in HeLa cells co-treated with LCFA or VLCFA (25 µM) in the presence of ELOVL1i (0.25 µM). Toxicity was normalized to the corresponding vehicle-treated controls. Data represent mean ± SD (n = 4). IC_50_ values were derived from non-linear regression with 95% confidence intervals.**d** Mitochondrial swelling in freshly isolated mouse liver mitochondria after direct addition of individual 1-deoxyDHCer, 1-deoxyCer, or canonical dihydroceramide (DHCer) species, as indicated.**e** Mitochondrial swelling in isolated liver mitochondria upon exposure to 1-deoxyDHCer (m18:0/24:1), in the presence or absence of the mitochondrial permeability transition pore (mPTP) inhibitor cyclosporin A (CsA, 10µM). **f** 1-deoxySa cytotoxicity in HeLa cells treated with CsA (10µM) or Bax inhibitor (Bax i, Bax inhibitor peptide V5, 10µM). Significance was assessed using two-tailed unpaired Student’s t-test with correction for multiple testing (two-stage step-up method of Benjamini, Krieger, and Yekutieli). *p < 0.05, **p < 0.01, ***p < 0.001.**g**, Schematic illustration of our proposed model of 1-deoxySL toxicity. We hypothesize that accumulation of very-long-chain (VLC) 1-deoxyDHCer species-particularly 1-deoxyDHCer (m18:0/24:1)- triggers opening of the mitochondrial permeability transition pore (mPTP), resulting in mitochondrial swelling, dysfunction, and BAX-mediated cell death.

Lipidomics profiling confirmed the selective increase in the corresponding LC or VLC 1-deoxyDHCer species following supplementation (Figure 5b). Toxicity assays revealed that primarily nervonate (24:1), which promoted the formation of 1-deoxyDHCer (m18:0/24:1) (also referred as nervonyl-1-deoxyDHCer), induced the strongest cytotoxic effect, followed by lignocerate (24:0) and palmitate (16:0) (Figure 5c). These findings identify nervonyl-1-deoxyDHCer (m18:0/24:1) as the most toxic 1-deoxySL species generated in the cells.

To evaluate the impact of these individual species on mitochondrial integrity, we conducted a mitochondrial swelling assay using freshly isolated mitochondria exposed directly to synthetic LC and VLC 1-deoxySL. Among the tested species, 1-deoxyDHCer (m18:0/24:1) induced the most signficant swelling reponse, followed by 1-deoxyDHCer (m18:0/24:0) and (m18:0/16:0), whereas unsaturated 1-deoxyCer (e.g., m18:1/16:0, m18:1/24:0) and canonical dihydroceramides (d18:0/xx:y) showed no effect (Figure 5d). This demonstrates that 1-deoxyDHCer (m18:0/24:1) directly perturbs mitochondrial integfrity.

Mitochondrial swelling is a hallmark of mitochondrial permeability transition pore (mPTP) opening, a well known trigger for apoptosis^27^. To test whether mPTP is involved in 1-deoxySL-induced toxicity, we treated isolated mitochondria with 1-deoxyDHCer (m18:0/24:1) in the presence or absence of the mPTP inhibitor cyclosporin A (CsA). CsA significantly decreased the mitochondrial swelling induced by this species (Figure 5e). Furthermore, in cellular toxicity assays, CsA rescued HeLa cells from 1-deoxySa-induced death (Figure 5f), implicating mPTP activation in the cytotoxic mechanism.

Since mPTP opening typically triggers a BAX-dependent apoptotic signaling, we next tested whether BAX inhibition can modulate 1-deoxySa toxicity^27^. The presence of a BAX inhibitor peptide (V5) significantly protected HeLa cells from 1-deoxySa-induced cell death (Figure 5f), indicating that BAX activation operates downstream of mPTP opening.

Taken together, these results demonstrate that 1-deoxyDHCer (m18:0/24:1) directly induces mitochondrial damage via mPTP opening and BAX activation, leading to cell death. This defines a novel molecular mechanism of 1-deoxySL cytotoxicity, with potential implications for the understanding of cellular basis of 1-deoxySL associated diseases (Figure 5g).

## Discussion

In this study, we identify a previously unrecognized molecular mechanism by which 1-deoxysphingolipids (1-deoxySLs) exert their cytotoxic effects. We demonstrated that mainly a single N-acylated 1-deoxyDHCeramide (m18:0/24:1) structure is responsible for toxicity and mitochondrial dysfunction associated with an opening of the mPTP and BAX-mediated cell death. This finding establishes a direct link between 1-deoxySL structure and a defined cell death pathway, providing new mechanistic insight into the pathophysiology of 1-deoxySL associated diseases.

1-deoxySLs are atypical sphingolipids generated via serine palmitoyltransferase (SPT) when alanine is utilized instead of serine. These lipids include long-chain bases (LCBs) such as 1-deoxysphinganine (1-deoxySa) and 1-deoxysphingosine (1-deoxySo), as well as their N-acylated derivatives (1-deoxyDHCer and 1-deoxyCer)^1,3^. The pathologically increased formation of 1-deoxySLs is implicated in a range of neurological conditions, including hereditary sensory and autonomic neuropathy type 1 (HSAN1), diabetic neuropathy, and macular telangiectasia (MacTel), as well as emerging roles in cancer metabolism and sarcopenia^5,9-13,17,28-31^.

While previous studies have shown that 1-deoxySLs are toxic in various cell types, especially neurons, the molecular basis of this toxicity has remained unclear^20,26,32^. Previous studies have shown that inhibiting ceramide synthase by fumonisin B1 (FB1) protects against the cytotoxic effects of 1-deoxysphingolipids. For example, FB1 was reported to rescue primary neurons from 1-deoxySL-induced degeneration and to restore fibroblast migration inhibited by 1-deoxySL accumulation^20,33^. Consistent with these findings, we observed that FB1 treatment significantly reduced the toxicity of 1-deoxySa. Mechanistically, FB1 inhibits the N-acylation of 1-deoxyLCBs, resulting in an accumulation of free 1-deoxySa and 1-deoxySo and a marked decrease in 1-deoxyDHCer and 1-deoxyCer levels. This supports the conclusion that N-acylated products, rather than the free LCBs, are responsible for toxicity.

In parallel, prior work has implicated fatty acid desaturase 3 (FADS3), an enzyme that introduces a Δ14(Z) double bond into sphingoid bases, as a modulator of 1-deoxySL toxicity. Specifically, FADS3 overexpression was shown to rescue cells from 1-deoxySa-induced toxicity, while FADS3 knockdown exacerbated toxicity, likely by favoring the accumulation of saturated, highly toxic 1-deoxyDHCer species^4^. In our study, we observed that 1-deoxySo, which is formed from 1-deoxySa by FADS3, is converted exclusively to 1-deoxyCer,and exhibited substantially lower toxicity than 1-deoxySa. These data, together with the protection by FB1, showed that 1-deoxyDHCer, but not 1-deoxySa or 1-deoxyCer, represents the principal cytotoxic metabolites within the 1-deoxySL pathway.

CRISPR-based genetic screens have emerged as a powerful and widely adopted tool in functional genomics ^34,35^. Unlike RNAi-based methods, CRISPRi achieves robust and specific gene silencing with reduced off-target effects, enabling the discovery of essential pathways and potential therapeutic targets in toxicity models. To systematically identify genetic regulators of 1-deoxySL toxicity, we employed a CRISPR interference (CRISPRi) screen in haploid K562 cells. Our screen identified five high-confidence hits—ELOVL1, HSD17B12, ACACA, PTPLB, and CERS2—all of which are involved in fatty acid elongation or ceramide biosynthesis. These enzymes regulate the formation and utilization of very-long-chain fatty acids (VLC-FAs), particularly 24-carbon saturated and monounsaturated species, which are subsequently incorporated by CERS2 into sphingolipids^36^. Notably, VLC-1-deoxyDHCer species such as m18:0/24:0 and m18:0/24:1 predominate in cells while canonical sphingolipids display a broader distribution of the N-acyl chain length (data not shown).

Functional validation using siRNA and CRISPR-Cas9 knockout models of ELOVL1 and CERS2 confirmed their essential role in mediating 1-deoxySL toxicity. Disruption of either enzyme significantly decreased VLC-1-deoxyDHCer formation and rescued cells from 1-deoxySa-induced toxicity, highlighting the critical importance of N-acyl chain length in determining cytotoxicity. These results are in agreement with prior yeast models showing a chain-length–dependent toxicity of 1-deoxySLs^37^.

To explore the therapeutic potential of this pathway, we employed a preclincally validated ELOVL1 inhibitor (22) that was originally developed in the context of X-linked adrenoleukodystrophy (X-ALD)^24,25^. Compound 22 has been shown to be pharmacologically active in mouse and rat models, with established preclinical safety and efficacy in cynomolgus monkeys^24,25^. Treatment with (22) blocked VLC-FA elongation and thereby suppressed the formation of toxic VLC-1-deoxyDHCer species. This resulted in a robust protection from 1-deoxySL-induced cytotoxicity in HeLa cells and neurotoxicity in primary dorsal root ganglion (DRG) neurons. Importantly, we validated the metabolic activity of (22) in vivo using a chicken embryo DRG flux assay, confirming its action in a neuronal context.

To further investigate the structure–toxicity relationship, we selectively stimulated the formation of individual 1-deoxyDHCer species in cells by supplementing defined LCFAs or VLCFAs in presence of (22) to avoid further metabolism of the added FAs by ELOVL1. Among all tested species, 1-deoxyDHCer (m18:0/24:1)—derived from nervonic acid (24:1)—was the most toxic, followed by m18:0/24:0 and m18:0/16:0. These findings point to nervonyl-1-deoxyDHCer (m18:0/24:1) as the main mediator of 1-deoxySL cytotoxicity. Of note, nervonic acid is enriched in myelin, suggesting a potential link between 1-deoxySL metabolism and demyelinating phenotypes observed in HSAN1 and related neuropathies.

Due to systemic mutations in SPT, 1-deoxySa are elevated in many tissues of HSAN1 patients. Nevertheless, the disease specifically affects sensory neuron function, leading to impaired pain and temperature sensation. Interestingly, previously reported single cell RNAseq data of neurons ^38^ indicate a co-expressing of ELOVL1 and CERS2 in specific nociceptor subtypes that also express also pain receptors such as TRPV1 or TRPA1 (Supplementary table 1). Based on our observations these neuronal subtypes are likely more susceptible to 1-deoxySL toxicity which might also explain the specificity of clinical symptoms seen in HSAN1 such as the loss of pain and temperature sensation. However, without further validation it is still speculative based and interventional studies are needed to confirm this association in more detail.

Previous reports have suggested that 1-deoxySLs localize to mitochondria, where they disrupt mitochondrial morphology^26^. Currently, it is not clear what exact biochemical or biophysical mechanisms relate to the differential toxicities of the individual 1-deoxySL species. For canonical Cer it has been shown that LC and VLC-Cer have opposing effects on Ca^2+^- induced mitochondrial swelling ^39^. Moreover, the N-acyl length and saturation of Cer has been shown to influence the biophysical properties of artificial membranes ^40^. We confirmed these findings using Seahorse experiments, fluorescence microscopy, and mitochondrial swelling assays. Supplementation with 1-deoxySa induces mitochondrial fragmentation, loss of respiratory and glycolytic capacity, and mitochondrial swelling. Notably, Nervonyl-1-deoxyDHCer (m18:0/24:1) caused the strongest mitochondrial swelling and cellular toxicity, which could be blocked by cyclosporin A (CsA), implicating mPTP opening as the underlying mechanism. Furthermore, inhibition of BAX, a downstream effector of mPTP-mediated apoptosis^27^, rescued 1-deoxySa induced cell death. These findings define a new mechanism of 1-deoxySL toxicity involving mitochondrial destabilization via mPTP and BAX activation.

Despite these insights, our study has certain limitations. The efficacy of (22) has not yet been tested in an animal model of 1-deoxySL associated diseases such as HSAN1 or diabetic neuropathy. Importantly, while (22) was designed to cross the blood–brain barrier for treating X-ALD^24,25^, this property may be less desirable for targeting peripheral neuropathies. Future work should therefore focus on developing ELOVL1 inhibitors optimized for peripheral selectivity and tested in PN relevant disease models.

In summary, our findings establish a direct link between the chemical structure of 1-deoxySLs and their cytotoxicity, revealing that both the saturation of the long-chain base and the length and unsaturation of the N-acyl chain determine toxicity. Most importantly, we identify 1-deoxyDHCer (m18:0/24:1) as a specific, mitotoxic species and uncover mPTP/BAX- associated mitochondrial dysfunction as a new molecular mechanism of 1-deoxySL-induced cell death. These findings support ELOVL1 inhibition as a promising therapeutic strategy for 1-deoxySL-associated diseases, including HSAN1, MacTel, and diabetic neuropathy.

## Methods

### Cell culture

HeLa and SH-SY5Y cell lines were grown in high-glucose Dulbecco’s Modified Eagle’s Medium (DMEM, Thermo Fisher Scientific) supplemented with 10% fetal bovine serum (FBS) and 1% Penicillin/Streptomycin (P/S). K562 human myeloid leukemia cells were cultured in RPMI1640 Medium GlutaMAX (Thermo Fisher Scientific) supplemented with 10 % FBS and 1% P/S. All cell lines were kept in the incubator at a 5% CO_2_ atmosphere and 37°C. Cells were tested for Mycoplasma contamination.

Differentiation of SH-SY5Y was performed in 96-well plates coated with freshly prepared laminin (24 hours, 10µg/mL). During the differentiation, cells were cultured for the first 7 days with media (DMEM) containing 1% P/S, 10% FBS and 10µM of Retinoic acid (R2625, Sigma-Aldrich). For the following 7 days of differentiation media (DMEM) containing 1% P/S, 10% FBS, 10µM of Retinoic acid and 100ng/mL Brain-derived neurotropic factor (BDNF, B3795, Sigma-Aldrich) was used.

HeLa CERS2 KO, CERS5/6 KO and ELOVL1 KO were prepared as described previously ^41^.

### Cultures of dissociated DRG neurons

Dissociated dorsal root ganglia (DRG) neurons were cultured as described previously^42^. Briefly, DRGs of 9-days old chicken embryos were dissected and collected in cold PBS. DRGs were centrifuges with 1000 rpm for 5 minutes at room temperature. PBS was exchanged with 0.25% Trypsin in PBS (15090-046, Invitrogen) containing DNase (final concentration: 0.2% ;10104159001, Roche) and incubated for 20 minutes at 37 °C. DRGs were collected by centrifugation, suspended in 1 ml DRG medium, and triturated with a fire-polished glass pipette with a diameter of about 0.5 mm. DRG media consisted of MEM with Glutamax (41090-028, Invitrogen), Albumax (4 mg/ml, 11020-021, Invitrogen), N3, and NGF (20 ng/ml, 13290-010, Invitrogen). Per well, 20000 cells were plated in 8-well Lab-Tek plates (177445, Nunc) and incubated at 37 °C with 5% CO_2_. After initial growth, 1-deoxySa and/or ELOVL1 inhibitor (compound 22 (22), HY-145272, MedChemExpress) were added to the corresponding wells. Cells were fixed after 40 hours of incubation with 4% PFA and washed 3x10 minutes with PBS.

### d3-1-deoxySa flux in chicken embryo derived dorsal root ganglia

To investigate whether ELOVL1 inhibition by (22) could influence the 1-deoxyDHCer profile from d3-1-deoxySa treatment, chicken embryos were utilized, as previously described^43,44^. Briefly, 4-days-old (E4) chicken embryos were windowed, and d3-1-deoxySa (50 µM, 50 µL) was injected to the embryo with or without (22) (2.5 µM, 50 µL). After 4 days, 7-days-old chicken DRGs were dissected according to previously established protocols^42^. For DRG dissection, embryos were pinned to a dissection plate, and the internal organs, ventral vertebrae, and spinal cord were carefully removed to expose the DRG. The DRGs were then collected in PBS, and then excess PBS was discarded. The tissues were subsequently frozen at -20° until lipid extraction and mass spectrometry analysis as described below.

### Silencing of ELOVL1

siRNAs targeting human ELOVL1 (s34994, Thermo Fisher Scientific) were used to silence (knockdown) ELOVL1 expression. ELOVL1 targetting siRNA or non-coding negative control Scramble (Scr, SR30004, OriGene) siRNAs were diluted to a final concentration 10 nM or 20 nM in reduced-serum media (Opti-MEM, Gibco). Transfection was performed using Lipofectamine RNAiMAX transfection Reagent (Thermo Fisher Scientific) according to the manufacturer’s recommendations. The media was replaced after 24 hours with fresh DMEM (10%FBS) and cells were grown in total for 72 hours before the start of labeling experiments. Knockdown efficiency was determined using qRT-PCR as described below.

### qRT-PCR

For qRT-PCR, cells were harvested using trypsinisation as described above. Cell pellets were lysed using TRIzol reagent and RNA was extracted using the phenol/chloroform method following manufacturer‘s instructions. RNA was purified using Ethanol and concentration/purity was determined using NanoDrop (Thermo Fisher Scientific). Reverse transcription was performed using Maxima Reverse Transcriptase following manufacturer‘s instructions (EP0742, Thermo Fisher Scientific).

qPCR reactions were performed using SYBR Green qPCR mastermix. Absolute concentration of mRNA was calculated from the dilution curve (1/10, 1/100, 1/1000 and 1/10000 dilution) of an adequate plasmid. GAPDH was used as an in-house gene loading control.

### Stable isotope labelling assay

For the SL labelling assay, cells were plated at 200 000 cells/mL in 6-well plates. Cells were grown for 48 hours to 70% confluency. 24 hours before harvesting, the medium was replaced with DMEM/10%FBS/1%P/S containing d_3_-1-deoxysphinganine (860474, Avanti Polar Lipids) and additional treatments (compound 22 (22), HY-145272, MedChemExpress; FB1, F1147, Sigma-Aldrich) as indicated in the figures. Cells were harvested by trypsinisation and counted using Beckman Coulter Z2 (Beckman Coulter). Next, cells were centrifuged at 850g at 4°C and cell pellets washed 2 times with cold PBS. Cell pellets were then frozen and kept at -20°C until further analysis.

### Fatty acids supplementation

Fatty acids were supplemented to cells as sodium salts conjugated to Bovine Serine Albumin (BSA). First, 10µmol of fatty acids were weighted into an Eppendorf tube and EtOH (100µL), H_2_O (300µL) and 0.1M NaOH (100µL) were added. Next, the mixture was heated at 75°C and mixed for 2 hours (Thermomixer (Eppendorf)). Afterwards, 500µL of H_2_O were added (yielding total 10mM concentration). BSA complexes were prepared by the addition of 100µL of fatty acid sodium salts to 900µL BSA (Essentially fatty acids free (Thermo Fisher Scientific)) in DMEM (6.66mg/mL) and shaking at 30°C for 1 hour ((Thermomixer (Eppendorf)) and used directly for isotope labelling as described above. For a vehicle control, no fatty acids were added and the whole process was performed accordingly.

### Lipid extraction and analysis

Lipidomics analysis was performed as described previously ^45^.Extraction was performed by mixing (Thermomixer (Eppendorf), 60 minutes, 37°C) the cell pellets, plasma (100µL), or tissue homogenate with extraction buffer consisting of a mixture: methanol: methyl *tert-*butyl ether: chloroform 4:3:3 (v/v/v) and internal standards. After centrifugation (16 100 rpm, 10 minutes, 37°C), the single-phase supernatant was collected, dried under N2, and stored at -20°C. Before analysis, lipids were dissolved in 100µL of MeOH Thermomixer (Eppendorf), 60 minutes, 37°C) and separated on a C30 Accucore LC column (Thermo Fisher Scientific, 150mm x 2.1 mm x 2.6 µm) or C18 ACQUITY UPLC CSH (Waters, 150mm x 2.1 mm x 1.7µm) using gradient elution with A) Acetonitrile: Water (6:4) with 10mM ammonium acetate and 0.1% formic acid and B) Isopropanol: Acetonitrile (9:1) with 10mM ammonium acetate and 0.1% formic acid at a flow rate of 260µL/minute using Transcend UHPLC pump (Thermo Fisher Scientific) at 50°C. Following gradient was used: 0min 30% (B), 0.5min 30% (B), 2min 43% (B), 3.3min 55% (B), 12min 75% (B), 18min 100% (B), 25min 100% (B), 25.5min 30% (B), 29.5min 30% (B). Eluted lipids were analysed by a Q-Exactive plus HRMS (Thermo Fisher Scientific) in positive and negative modes using heated electrospray ionisation (HESI, Shealth gas flow rate=40, Aux gas flow rate=10, Sweep gas flow rate=0, Spray voltage=3.50, Capillary temperature=320°C, S-lens RF level=50.0, Aux gas heater temperature=325°C). MS2 fragmentation spectra were recorded in data-dependent acquisition mode with top10 approach and constant collision energy 25eV. 140 000 resolution was used for full MS1 and 17 500 for MS2. Peak integration was performed with TraceFinder 4.1, Skyline daily. Lipids were identified by predicted mass (resolution 5ppm), retention time (RT) and specific fragmentation patterns using in house made, Lipidcreator and MSDIAL compound databases. Isotopic enrichment was tracked by M+3 for d_3_-1-deoxySa treated cells. Next, lipid concentrations were normalized to the corresponding internal standards (one per class) and cell number.

#### List of internal standards

d5-1-deoxymethylsphinganine (m17:0, 860476, Avanti Polar Lipids) 100pmol/sample

1-deoxyDHCeramide (m18:0/12:0, 860460P, Avanti Polar Lipids) 100pmol/sample

1-deoxyCeramide (m18:1/12:0, 860455, Avanti Polar Lipids) 100pmol/sample

DHCeramide (d18:0/12:0, 860635 Avanti Polar Lipids) 100pmol/sample

Ceramide (d18:1/12:0, 860512, Avanti Polar Lipids) 100pmol/sample

SM (d18:1/12:0, 860583, Avanti Polar Lipids) 100pmol/sample

Glucosylceramide (d18:1/8:0, 860540, Avanti Polar Lipids) 100pmol/sample

SPLASH 2.5µL/sample (330707, Avanti Polar Lipids)

Transitions used for the identification of Sphingolipids and 1-deoxySphingolipids: 1-deoxyCeramides and 1-deoxyDHCeramides:

[M+H]^+^ → [M+H – H_2_O]^+^, [M+H]^+^ → [M+H – H_2_O - FA]^+^

Ceramides and dihydroCeramides:

[M+H]^+^ → [M+H - H_2_O]^+^, [M+H]^+^ → [M+H - H_2_O - FA]^+^, [M+H]^+^ → [M+H - 2xH_2_O - FA]^+^

Free atypical long-chain bases

[M+H]^+^ → [M+H - H_2_O]^+^

Note: FA represents corresponding fatty acyl.

### Toxicity assays

For the toxicity assay cells were grown for 72 hours in the 96-well plates in DMEM (supplemented with 10% FBS and 1% P/S) with added treatments as indicated in the figures. All conditions were corrected for the solvent concentration. Chemicals used were dissolved in DMSO to 1mM concentration and added directly to the media.

Chemicals used: Fuminosin B1 (F1147, Sigma-Aldrich); Compound 22 (HY-145122, MedChemExpress); Cyclosporin A (HY-B0579, MedChemExpress); Bax inhibitor peptide V5 (HY-P0081, MedChemExpress), 1-deoxySa (860493, Avanti Polar Lipids).

The number of viable cells was determined by quantitation of the ATP present using CellTiter-Glo **®** Luminescent Cell Viability Assay (G7570, Promega) according to the manufacturer‘s recommendations. Chemiluminesce signal was collected using TECAN infinite M 200 Pro reader. Data were normalized to the average of Vehicle treated wells.

### Immunocytochemistry of DRG dissociated sensory neurons

Cells were permeabilized with 0.1% Triton-X 100 in PBS and blocked for 15 minutes with 5% FCS in PBS (blocking buffer) at room temperature. Primary antibodies were diluted in blocking buffer and added to the cells for 1 hour. Primary antibodies included anti-Neurofilament-M (1:2000; clone RMO270, RRID: AB_2315286, Invitrogen) and anti-TOMM20 (rabbit anti-TOMM20, HPA011562, 1:800, Sigma-Aldrich). Cells were washed with PBS and incubated with secondary antibodies diluted in blocking buffer for another hour. Then cells were stained with Hoechst (2.5 µg/ml in PBS, H3570) for 5 minutes and rinsed three times with PBS. Finally, cells were mounted with Mowiol/Dabco.

### Microscopy and quantification of neurite density

Neurons were imaged with a 20x air objective and an Olympus BX63 upright microscope. Neurite growth was analyzed with Fiji/ImageJ. For this purpose, a region of interest (6500x6500 micrometers) was chosen in the center of each well. Images were transformed into 8-bit and a threshold was applied. The total area of neurons and neurites was assessed with the “Measure” feature.

### Live cell microscopy

For the live cell microscopy cells were grown in 96-well plates (Ibidi) and the indicated treatment was added by exchange of the media. Then, the plate was immediately transferred to a live-cell imaging microscope (Olympus IX81) with a motorized stage, fitted with an incubator with pre-heated humidified atmosphere (Ibidi mixer). Phase-contrast images were acquired at 20x magnification every hour for 48 hours. At all procedures cells were kept at 5% CO_2_ and 37°C.

### Whole-genome CRISPRi 1-deoxySa toxicity screen

CRISPRi toxicity screen was performed using the K562 cell line as described in previously published screens ^46,47^. Briefly, K562 cells stably expressing dCas9-KRAB were transduced with the CRISPRi v2 sgRNA library. This library contains 10 sgRNAs for the ∼20K protein coding genes, along with 10 computer generated, non-targeting sgRNAs clustered into 15K “pseudogenes”. In all 350K sgRNAs were used with a 60% targeting, 40% non-targeting ratio. 48 hours later, cells were selected with 0.75 µg/ml puromycin for 72 hours. Next, cells were recovered from the selection with puromycin-free media for 48 hours. Then, two replicates of 200M cells were split into two groups: one group was not treated, while the other was treated with 1.5µM 1-deoxySa (860493, Avanti Polar Lipids). For each flask in the untreated and treated groups, the cells were kept to 0.5 million cells/ml daily. This means 1.5µM 1-deoxySa caused a 50% growth inhibition. Treatment was continued until untreated cells doubled five to eight more times than the treated cells. Genomic DNA was collected from all harvested samples. The genomic regions containing the inserted sgRNAs were amplified by PCR, and abundances were determined by next-generation sequencing. The analysis protocol including code is available here: https://github.com/mhorlbeck/ScreenProcessing.

### Mitochondria isolation from liver

Functional mitochondria were isolated from the livers of 10-12 week-old female C57/BJ mice (modification of the method described in ^48^. Briefly, after excision, the livers were placed in an ice-cold isolation buffer (200 mM mannitol, 50 mM sucrose, 5 mM KH_2_PO_4_, 5 mM MOPS, 0.1% fatty acid free BSA, 1 mM EGTA, adjusted to pH 7.15 with KOH), homogenized with a motor-driven tightly fitting glass-Teflon Potter grinder operated at 1 600 rpm and centrifuged at 3 minutes (4°C, 1 100g). The supernatant was centrifuged at 4°C for 10 minutes at 10,000g. The pellet containing the mitochondrial fraction was resuspended in 200-400 µl of assay buffer (120 mM KCl, 10 mM Tris, 5 mM KH_2_PO_4_, pH 7.4) and centrifuged for 10 minutes (4°C, 10 000g). Supernatant was decanted and the pellet was resuspended in the assay buffer. Mitochondria were kept on ice and an aliquot was used for the assessment of protein content by BCA assay (Thermo Fisher Scientific) according to the manufacturer‘s recommendations.

### Mitochondrial swelling assay

The activation of the mitochondrial permeability transition pore causes mitochondrial swelling, which is measured spectrophotometrically as a decrease in absorbance at 540 nm as described previously^39^. Briefly, the activation of the mitochondrial permeability transition pore causes mitochondrial swelling, which is measured spectrophotometrically as a decrease in absorbance at 540 nm. 250 µg aliquots of mitochondria suspension were distributed in a clear bottom 96-well plate, and then incubated directly with the indicated 1-deoxyDHCer, 1-deoxyCer or DHCer (Ethanolic solutions) at 4x10-3 nmol per ug of protein^39^. After 1 hour, the decline in the absorbance at 540 nm was measured. For the inhibition of mPTP, cyclosporine A (CsA, 10µM) was used. CaCl2 (Ca2+, 500 μM) was used as the positive control for the experiments. For the time series of measurement, the absorbance at 540 nm was recorded every 10 min after the indicated treatments until 1 hour.

### Seahorse assay

The Seahorse extracellular flux (XF) 24 analyzer (Agilent) was employed to analyze cellular oxygen consumption rate (OCR) and extracellular acidification rate (ECAR). Initially, 20000 cells were seeded onto a Seahorse microplate containing 250 μL of growth medium containing treatment as indicated in the figures. 72 hours later, Seahorse XF basal medium was added (maintained at pH 7.4). The medium was maintained in a CO_2_-free incubator at 37°C for an hour. The subsequent mitochondrial stress test or glycolysis stress test was executed to quantify OCR and ECAR, respectively.

For the mitochondrial stress test, the XF basal medium was enriched with 10 mM glucose, 1 mM pyruvate, and 2 mM glutamine. OCR and ECAR measurements were taken during distinct phases: baseline, after adding oligomycin (final concentration 1 μM), following the introduction of carbonyl cyanide-p-trifluoromethoxyphenyl-hydrazone (FCCP; final concentration 1 μM), and subsequent to the mixture of rotenone/antimycin A addition (final concentration of 0.5 μM each).

Conversely, the glycolysis stress test was conducted using XF basal medium supplemented with 2 mM glutamine. OCR and ECAR observations were collected under the subsequent conditions: baseline, after adding glucose (final concentration 10 mM), following the introduction of oligomycin (final concentration 1 μM), and after adding 2-deoxyglucose (2-DG; final concentration 50 mM).

The data were analysed using Agilent’s Wave software. Lastly, flux rates were normalized to Crystal violet signal (measured by Tecan reader, 565nm) after fixation with 4% paraformaldehyde (PFA, Thermo Fisher Scientific).

### Gene expression analysis in dorsal root ganglion (DRG) cells

Gene expression data in individual DRG cells, obtained from single cell RNA-seq data originally published by Usoskin et al, was downloaded from http://linnarssonlab.org/drg/. The expression of each gene was illustrated as reads-per-million (RPM), and the maximal expression level for each gene of interest was calculated by averaging the three highest values across all analyzed cells. A cell expressing the gene at levels higher than 5% of the maximal level was considered positive for the gene, which is a similar criterion as Usoskin et al, with the exception that we also included non-neuronal cell types in our analysis. Within the 11 neuronal cell types and non-neuronal cells, we calculated the fraction of cells expressing each gene of interest. In addition, we calculated the fraction of cells positive for both ELOVL1 and CERS2 in each cell type.

### Data analysis, statistics, writing and figures

All the data analysis and figure preparations were performed using GraphPad Prism 9.5.1 and Excel. Statistical analysis was performed using GraphPad Prism 9.5.1. Gene set enrichment analysis was performed using WEB-based Gene SeT AnaLysis Toolkit (WebGestalt.org). Illustrations were made using Biorender webpage. Chemical structures were drawn using Chemdraw Ultra 12.0. ChatGPT 4o (OpenAI) was used to improve grammar and language clarity; no text or data were generated by the model. Final figures were prepared using Affinity Designer 2.

## Author contributions

MA performed all the experiments and analysis, interpreted the data and wrote the manuscript, KG supervised the study and interpreted the data, EY, MS, ES, RD performed the DRG sensory neurons preparations and analysis, LJ and PP performed CRISPRi screen, SM differentiated SH-SY5Y neurons, HaT generated CERS2, CERS5/6 KO and ELOVL1 cell lines, HP prepared fatty acid complexes, GZ performed mitochondrial swelling experiments, HT supervised the study and revised the manuscript.

## Acknowledgements

This project was supported by the Swiss National Science Foundation (SNF 310030_215134) and by the SNF under the frame of the European Joint Program on Rare Diseases (EJP RD+SNF 32ER30_187505) and COST Action 19105-Pan-European Network in Lipidomics and EpiLipidomics (EpiLipidNET). T.Ha. was supported by the NCCR Chemical Biology (185898 attributed to Howard Riezman). Additionally, T.Ha. was supported by the French Government (National Research Agency, ANR) through the "Investments for the Future" programs LABEX SIGNALIFE ANR-11-LABX-0028 and IDEX UCAJedi ANR-15-IDEX-01, the ATIP-Avenir program (CNRS/Inserm). We also would like to acknowledge Howard Riezman for his support on this project and Aaliyah Chaussin for her contribution in making the KO cell lines.

---

**Supplemental Table 1.**
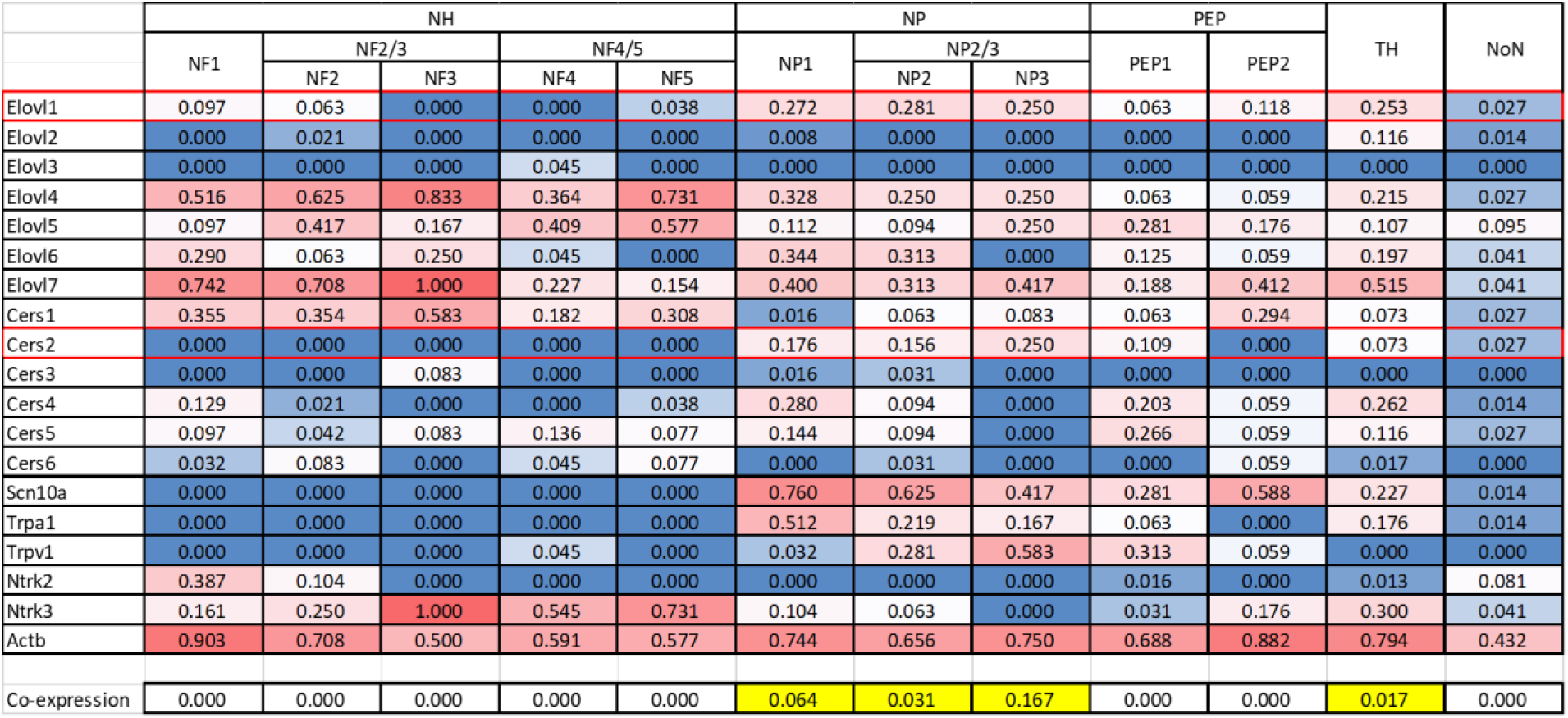
Gene expression profiles in dorsal root ganglion cells. CERS2 and ELOVL1 are co-expressed in nociceptors related to pain and temperature perception. DRG neurons classified based on single cell RNA-seq can be divided into 11 types (NF1-5, NP1-3, PEP1-2, and TH) 28. A co-expression of ELOVL1 and CERS2 is primarily seen in the neuron classes NP1-3, which express pain receptors (Scn10a, Trpa1, Trpv1) and in the TH subtype, which are involved in mechanical pain. No strong ELOVL1 and CERS2 co-expression is seen in other neuron types such as low threshold mechanoreceptors or proprioceptive neurons, nor in non-neuronal cell types. Other ELOVL and CERS isoforms are either ubiquitously expressed in all neuron subtypes, or not expressed at all. Thus, the distinctive ELOVL1 and CerS2 co-expression in a specific subset of DRG neurons potentially contributes to tissue selectivity.

## References

1 Hannun, Y. A. & Obeid, L. M. Sphingolipids and their metabolism in physiology and disease. Nat Rev Mol Cell Bio 19, 175–191 (2018). 10.1038/nrm.2017.107

2 Penno, A. et al. Hereditary Sensory Neuropathy Type 1 Is Caused by the Accumulation of Two Neurotoxic Sphingolipids. Journal of Biological Chemistry 285, 11178–11187 (2010). 10.1074/jbc.M109.092973

3 Lone, M. A., Santos, T., Alecu, I., Silva, L. C. & Hornemann, T. 1-Deoxysphingolipids. Biochim Biophys Acta Mol Cell Biol Lipids 1864, 512–521 (2019). 10.1016/j.bbalip.2018.12.013

4 Karsai, G. et al. FADS3 is a Delta14Z sphingoid base desaturase that contributes to gender differences in the human plasma sphingolipidome. J Biol Chem 295, 1889–1897 (2020). 10.1074/jbc.AC119.011883

5 Cordes, T. et al. 1-Deoxysphingolipid synthesis compromises anchorage-independent growth and plasma membrane endocytosis in cancer cells. J Lipid Res 63, 100281 (2022). 10.1016/j.jlr.2022.100281

6 Rosarda, J. D. et al. Imbalanced unfolded protein response signaling contributes to 1-deoxysphingolipid retinal toxicity. Nat Commun 14, 4119 (2023). 10.1038/s41467-023-39775-w

7 Sanchez, A. M. et al. Spisulosine (ES-285) induces prostate tumor PC-3 and LNCaP cell death by de novo synthesis of ceramide and PKC zeta activation. European Journal of Pharmacology 584, 237–245 (2008). 10.1016/j.ejphar.2008.02.011

8 Cuadros, R. et al. The marine compound spisulosine, an inhibitor of cell proliferation, promotes the disassembly of actin stress fibers. Cancer Lett 152, 23–29 (2000).

9 Rotthier, A. et al. Characterization of two mutations in the SPTLC1 subunit of serine palmitoyltransferase associated with hereditary sensory and autonomic neuropathy type I. Hum Mutat 32, E2211–2225 (2011). 10.1002/humu.21481

10 Rotthier, A. et al. Mutations in the SPTLC2 subunit of serine palmitoyltransferase cause hereditary sensory and autonomic neuropathy type I. Am J Hum Genet 87, 513–522 (2010). https://doi.org/S0002-9297(10)00478-7 [pii] 10.1016/j.ajhg.2010.09.010

11 Suriyanarayanan, S. et al. The Variant p.(Arg183Trp) in SPTLC2 Causes Late-Onset Hereditary Sensory Neuropathy. Neuromolecular Med 18, 81–90 (2016). 10.1007/s12017-015-8379-1

12 Murphy, S. M. et al. Hereditary sensory and autonomic neuropathy type 1 (HSANI) caused by a novel mutation in SPTLC2. Neurology 80, 2106–2111 (2013). 10.1212/WNL.0b013e318295d789

13 Othman, A. et al. Plasma 1-deoxysphingolipids are predictive biomarkers for type 2 diabetes mellitus. BMJ open diabetes research & care 3, e000073 (2015). 10.1136/bmjdrc-2014-000073

14 Othman, A. et al. Lowering plasma 1-deoxysphingolipids improves neuropathy in diabetic rats. Diabetes 64, 1035–1045 (2015). 10.2337/db14-1325

15 Hornemann, T. Serine deficiency causes complications in diabetes. Nature (2023). 10.1038/d41586-023-00054-9

16 Handzlik, M. K. et al. Insulin-regulated serine and lipid metabolism drive peripheral neuropathy. Nature (2023). 10.1038/s41586-022-05637-6

17 Gantner, M. L. et al. Serine and Lipid Metabolism in Macular Disease and Peripheral Neuropathy. N Engl J Med 381, 1422–1433 (2019). 10.1056/NEJMoa1815111

18 Esaki, K. et al. L-Serine Deficiency Elicits Intracellular Accumulation of Cytotoxic Deoxysphingolipids and Lipid Body Formation. J Biol Chem 290, 14595–14609 (2015). 10.1074/jbc.M114.603860

19 Alecu, I. et al. Localization of 1-deoxysphingolipids to mitochondria induces mitochondrial dysfunction. Journal of Lipid Research 58, 42–59 (2017). 10.1194/jlr.M068676

20 Lauterbach, M. A. et al. 1-Deoxysphingolipids cause autophagosome and lysosome accumulation and trigger NLRP3 inflammasome activation. Autophagy 17, 1947–1961 (2021). 10.1080/15548627.2020.1804677

21 Wilson, E. R. et al. Hereditary sensory neuropathy type 1-associated deoxysphingolipids cause neurotoxicity, acute calcium handling abnormalities and mitochondrial dysfunction in vitro. Neurobiol Dis 117, 1–14 (2018). 10.1016/j.nbd.2018.05.008

22 Karsai, G. et al. Metabolism of HSAN1- and T2DM-associated 1-deoxy-sphingolipids inhibits the migration of fibroblasts. J Lipid Res 62, 100122 (2021). 10.1016/j.jlr.2021.100122

23 Ohno, Y. et al. ELOVL1 production of C24 acyl-CoAs is linked to C24 sphingolipid synthesis. Proc Natl Acad Sci U S A 107, 18439–18444 (2010). 10.1073/pnas.1005572107

24 Come, J. H. et al. Discovery and Optimization of Pyrazole Amides as Inhibitors of ELOVL1. J Med Chem 64, 17753–17776 (2021). 10.1021/acs.jmedchem.1c00944

25 Boyd, M. J. et al. Discovery of Novel, Orally Bioavailable Pyrimidine Ether-Based Inhibitors of ELOVL1. J Med Chem 64, 17777–17794 (2021). 10.1021/acs.jmedchem.1c00948

26 Alecu, I. et al. Localization of 1-deoxysphingolipids to mitochondria induces mitochondrial dysfunction. J Lipid Res 58, 42–59 (2017). 10.1194/jlr.M068676

27 Bernardi, P. et al. Identity, structure, and function of the mitochondrial permeability transition pore: controversies, consensus, recent advances, and future directions. Cell Death Differ 30, 1869–1885 (2023). 10.1038/s41418-023-01187-0

28 Fridman, V. et al. Altered plasma serine and 1-deoxydihydroceramide profiles are associated with diabetic neuropathy in type 2 diabetes and obesity. J Diabetes Complications 35, 107852 (2021). 10.1016/j.jdiacomp.2021.107852

29 Dohrn, M. F. et al. Elevation of plasma 1-deoxy-sphingolipids in type 2 diabetes mellitus: a susceptibility to neuropathy? European Journal of Neurology 22, 806–e855 (2015). 10.1111/ene.12663

30 Green, C. R. et al. Divergent amino acid and sphingolipid metabolism in patients with inherited neuro-retinal disease. Mol Metab 72, 101716 (2023). 10.1016/j.molmet.2023.101716

31 Laurila, P. P. et al. Sphingolipids accumulate in aged muscle, and their reduction counteracts sarcopenia. Nat Aging 2, 1159–1175 (2022). 10.1038/s43587-022-00309-6

32 Guntert, T. et al. 1-Deoxysphingolipid-induced neurotoxicity involves N-methyl-d-aspartate receptor signaling. Neuropharmacology 110, 211–222 (2016). 10.1016/j.neuropharm.2016.03.033

33 Karsai, G. et al. Metabolism of HSAN1-and T2DM-associated 1-deoxy-sphingolipids inhibits the migration of fibroblasts. Journal of Lipid Research 62 (2021). https://doi.org/ARTN 100122 10.1016/j.jlr.2021.100122

34 Horlbeck, M. A. et al. Compact and highly active next-generation libraries for CRISPR-mediated gene repression and activation. Elife 5 (2016). 10.7554/eLife.19760

35 Gilbert, L. A. et al. Genome-Scale CRISPR-Mediated Control of Gene Repression and Activation. Cell 159, 647–661 (2014). 10.1016/j.cell.2014.09.029

36 Siddiqui, A. J. et al. Therapeutic Role of ELOVL in Neurological Diseases. ACS Omega 8, 9764–9774 (2023). 10.1021/acsomega.3c00056

37 Haribowo, A. G. et al. Cytotoxicity of 1-deoxysphingolipid unraveled by genome-wide genetic screens and lipidomics in Saccharomyces cerevisiae. Mol Biol Cell 30, 2814–2826 (2019). 10.1091/mbc.E19-07-0364

38 Usoskin, D. et al. Unbiased classification of sensory neuron types by large-scale single-cell RNA sequencing. Nat Neurosci 18, 145–153 (2015). 10.1038/nn.3881

39 Novgorodov, S. A., Gudz, T. I. & Obeid, L. M. Long-chain ceramide is a potent inhibitor of the mitochondrial permeability transition pore. Journal of Biological Chemistry 283, 24707–24717 (2008). 10.1074/jbc.M801810200

40 Pinto, S. N., Silva, L. C., Futerman, A. H. & Prieto, M. Effect of ceramide structure on membrane biophysical properties: the role of acyl chain length and unsaturation. Biochim Biophys Acta 1808, 2753–2760 (2011). 10.1016/j.bbamem.2011.07.023

41 Harayama, T., et al. Establishment of a highly efficient gene disruption strategy to analyze and manipulate lipid co-regulatory networks. bioRxiv, 2020: p. 2020.11.24.395632. (2020).

42 Yusifov, E., Dumoulin, A. & Stoeckli, E. T. Investigating Primary Cilia during Peripheral Nervous System Formation. Int J Mol Sci 22 (2021). 10.3390/ijms22063176

43 Wei, L. et al. Rho kinases play an obligatory role in vertebrate embryonic organogenesis. Development 128, 2953–2962 (2001).

44 Duess, J. W., Fujiwara, N., Corcionivoschi, N., Puri, P. & Thompson, J. ROCK inhibitor (Y-27632) disrupts somitogenesis in chick embryos. Pediatr Surg Int 29, 13–18 (2013). 10.1007/s00383-012-3202-7

45 Lone, M. A. et al. SPTLC1 variants associated with ALS produce distinct sphingolipid signatures through impaired interaction with ORMDL proteins. J Clin Invest 132 (2022). 10.1172/JCI161908

46 Horlbeck, M. A. et al. Mapping the Genetic Landscape of Human Cells. Cell 174, 953–967 e922 (2018). 10.1016/j.cell.2018.06.010

47 Yu, Z. et al. Identification of a transporter complex responsible for the cytosolic entry of nitrogen-containing bisphosphonates. Elife 7 (2018). 10.7554/eLife.36620

48 Frezza, C., Cipolat, S. & Scorrano, L. Organelle isolation: functional mitochondria from mouse liver, muscle and cultured fibroblasts. Nat Protoc 2, 287–295 (2007). 10.1038/nprot.2006.478

